# Experimental pieces of evidence for *Mycobacterium ulcerans* dormancy

**DOI:** 10.1101/2020.08.27.269712

**Authors:** A. Loukil, R. Lalaoui, H. Bogreau, S. Regoui, M. Drancourt, N Hammoudi

## Abstract

**Background:** Whether *Mycobacterium ulcerans*, the etiological agent of the neglected Buruli ulcer in numerous tropical countries, would exist in a dormant state as reported for closely related *Mycobacterium* species, is not established.

**Methodology:** Six *M. ulcerans* strains were exposed to a progressive depletion in oxygen for two months, using a previously described Wayne model of dormancy; and further examined by microscopy using DDD staining, microcalorimetry and subculture in the presence of dead and replicative *M. ulcerans* as controls.

**Principal Findings/Conclusions:** *M. ulcerans* CU001 strain died during the progressive oxygen depletion and four of five remaining strains exhibited Nile Red-stained intracellular lipid droplets after DDD staining and a 14-20-day regrowth when exposed to ambient air, diagnosing dormancy. A fifth *M. ulcerans* 19423 strain stained negative in DDD and slowly regrew in 27 days. Three tested *M. ulcerans* strains yielded microcalorimetric pattern similar to that of the negative (dead) homologous controls, differing from that of the homologous positive (replicative) controls. The relevance of these experimental observations, suggesting a previously unreported dormancy state of *M. ulcerans*, needs to be investigated in the natural ecological niches where *M. ulcerans* thrive and in Buruli ulcer lesions.

**Author summary:** *Mycobacterium ulcerans* is an environmental opportunistic pathogen of mammals and humans, causing a subcutaneous necrotizing infection named Buruli ulcer. Molecular detection of *M. ulcerans* DNA revealed different ecological niches where *M. ulcerans* may thrive, but the molecular biology approach does not catch the physiological state of *M. ulcerans* in these different ecological niches. Thus, the reservoir and the mode of transmission of *M. ulcerans* remain elusive. Here, we investigated experimental dormancy of *M. ulcerans* by using a previously described Wayne model of dormancy coupled with microscopy using DDD staining, microcalorimetry and subculture. Our findings demonstrate for the first time that some *M. ulcerans* strains exhibit a physiological state of dormancy; potentially limiting isolation and culture of *M. ulcerans* from environmental niches.

## INTRODUCTION

*Mycobacterium ulcerans* is an environmental non-tuberculous mycobacterium acting as an opportunistic pathogen of mammals and humans in whom it is responsible for a subcutaneous necrotizing infection named Buruli ulcer [1]. Buruli ulcer lesions are primarily caused by cytotoxic macrolide exotoxins named mycolactones, naturally secreted by *M. ulcerans*, of which seven different natural variants have been characterized in the several genotypes of *M. ulcerans* clinical isolates [2, 3]. All these variants target the Wiskott-Aldrich protein Arp2/3 in targeted cells, yet exhibiting various degrees of cytotoxicity, *in vitro* [4, 5]. These observations all issued from the investigations of *M. ulcerans* clinical isolates as no isolate has been made from the environmental sources where this opportunistic pathogen is regularly detected, except from one firmly documented *M. ulcerans* isolate made from one *Gerris* sp. aquatic insect from a Buruli ulcer-endemic area in Benin, West Africa [6]; contrasting with numerous *M. ulcerans* isolates made from clinical sources in Buruli ulcer patients and some mammals [7, 8].

The sharp discrepancy in the yield of *M. ulcerans* recovered from clinical sources versus environmental sources has been attributed to the intrinsic fastidiousness of the pathogen which is requiring a narrow 28-30°C growth temperature and specific culture medium; and to the overgrowth of reportedly contaminating bacteria and fungi [9]. Yet, none of these two limitations really differentiate clinical from environmental samples so that the exact reason for the tremendous difficulties in isolating environmental *M. ulcerans* remained largely unknown.

We herein explored experimentally a third line of explanation by investigating the possibility that *M. ulcerans* may exhibit a dormant state, less prone to laboratory cultivability than the dynamic state encountered in clinical samples. We indeed report that *M. ulcerans* may behave as a dormant mycobacterium and we showed that it is possible to convert the dormant into a dynamic state, opening a new venue for the isolation of still unexplored, environmental *M. ulcerans*.

## MATERIALS AND METHODS

### Wayne model of dormancy for *M. ulcerans*

Six different strains of *M. ulcerans* including *M. ulcerans* CU001, *M. ulcerans* ATCC 19423, *M. ulcerans* ATCC 33728, *M. ulcerans* ATCC 25900, *M. ulcerans* ATCC 25894 and *M. ulcerans* ATCC 25898 were investigated in this work. These strains are representative of the 02 genomic lineages currently described in *M. ulcerans* [10]; and originated from Ghana, Australia, Japan, USA, Ouganda, Benin, respectively [11]. These six *M. ulcerans* strains were cultured in Middlebrook 7H9 broth supplemented with 10% oleic acid/albumin/dextrose/catalase (OADC) (Becton Dickinson, Sparks, MD, USA) and 0.5% glycerol at 30°C in a shaking incubator. One non-inoculated tube was run in parallel as a negative control tube. These tubes were continuously incubated in the same conditions without changing the medium. After 50-day incubation of these tubes in this condition, the optical density (OD) of 0.2-0.3 at 600 nm., a 100-μL aliquot of each culture was suspended in one Hungate tube containing of 10 mL of Dubos broth (Becton Dickinson) in the presence of 1.5 μg/mL methylene blue (Sigma-Aldrich, Saint-Quentin-Fancy, France) as an indicator of oxygen depletion as previously described [12]. This Wayne model of dormancy initially described for *Mycobacterium tuberculosis* H37Rv relies on the gradual depletion of oxygen [13]. Accordingly, Hungate tubes were incubated at 30°C for up to 60 days with continuous shaking, with a weekly visual inspection (Fig. 1).

**Fig. 1.**
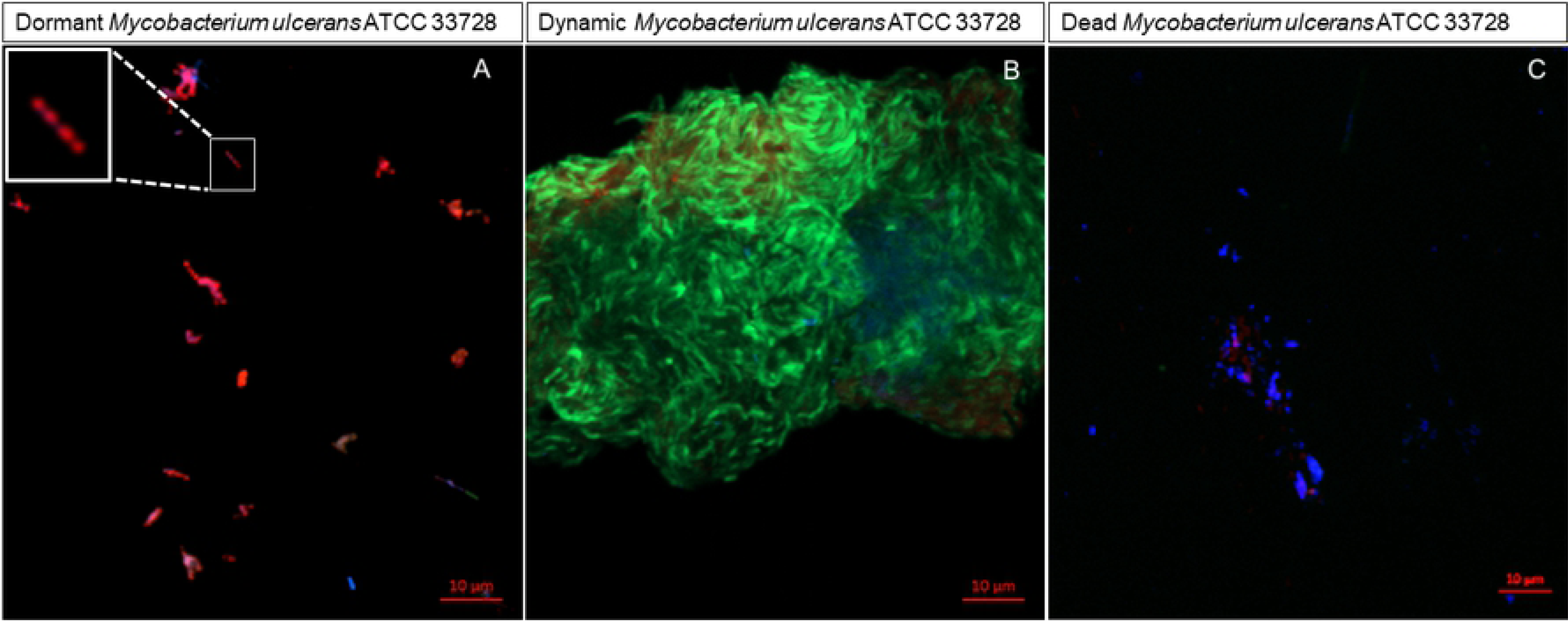
DDD staining of *M. ulcerans* ATCC 33728. A) Dormant *M. ulcerans* ATCC 33728 incubated for two months using the Wayne model of dormancy B) Dynamic *M. ulcerans* ATCC 33728 incubated in ambient atmosphere. C) Dead, heat-inactivated *M. ulcerans* ATCC 33728. Microscopic observations were performed using a 63X/1.4NA oil immersion objective of Zeiss spinning disk.

### DDD staining

After two-month incubation of the six *M. ulcerans* strains in the Hungate tubes in anaerobic conditions, a 20 μL-volume of each suspension was smeared on glass slide, air dried and stained using DDD staining protocol as previously described [12,13]. DDD staining was also applied on the aerobic and heat inactivated cultures of the six *M. ulcerans* strains. Microscopic observation was performed using a 63X/1.4NA oil immersion objective of Zeiss Spinning Disk Confocal microscopy driven by Zen software (Zeiss, Marly-le-Roi, France).

### Microcalorimetry test

Isothermal microcalorimetry was used to evaluate heat production by three *M. ulcerans* strains (*M. ulcerans* 33728, *M. ulcerans* 25900 and *M. ulcerans* 25894 strains) using a CalScreener prototype microcalorimeter (Symcel, Solna, Sweden). Briefly, 10^6^ colony forming units (CFUs) / mL of Middlebrook 7H9 medium supplemented with 10% OADC of dynamic mycobacteria of each one of the three investigated strains, were incubated for 1h at 100°C and further sonicated at 37 kHz for 30 minutes in order to get so-called dead mycobacteria. Then, 10^6^ CFUs / mL dead and 10^6^ CFUs / mL dynamic mycobacteria (in Middlebrook medium) and 10^6^ CFUs / mL dormant mycobacteria (in Dubos medium) were incubated at 30°C for 17 hours before being prepared for microcalorimetry. For microcalorimetry, 200 μL of each suspension were inoculated into 400-μL titanium tubes in triplicate, 09 tubes containing 200 μL of Middlebrook 7H9 medium supplemented with 10% OADC and 06 tubes containing 200 μL of Dubos broth were used as negative controls. These 24 sealed titanium tubes were placed in the microcalorimeter previously equilibrated at 30°C for 4 days. Individual heat production rates (i.e., W = J/s = heat flow or thermal power) of each titanium tube was measured continuously in real time during 100 hours as previously described [14].

### Subculture of dormant *M. ulcerans* strains in aerobic conditions

After the six *M. ulcerans* strains here investigated have been incubated for 60 days in anaerobic conditions in DUBOS medium, 100 μL were collected using 30-Gauge syringes to avoid oxygen penetration and centrifuged at 3,000 g for 15 min. The pellet was then seeded on Middlebrook 7H10 agar plates (Becton Dickinson) incubated at 30°C for 27 days in aerobic conditions. An identification of the cultured bacteria was carried out by molecular biology using real-time PCR targeting the ketoreductase b (kr-b) gene as previously described [15].

## RESULTS

### Visual inspection of tubes

After 24 to 44-day incubation at 30°C, all the *M. ulcerans*-inoculated tubes decolorized and a bacterial pellet appeared at the bottom except for three tubes inoculated with *M. ulcerans* ATCC CU001 strain; whereas the negative control tube remained blue-colored without any visible pellet (Fig. S1). All colonies were confirmed to be *M. ulcerans* (Supplementary Table).

### DDD staining result

After 2-month incubation in anaerobic conditions, DDD staining demonstrated the presence of red fluorescent-intracellular inclusions in four *M. ulcerans* strains except for *M. ulcerans* ATCC CU001 and *M. ulcerans* ATCC 19423 (Fig. 1, Fig. S2). These Nile red-positive lipidic bodies were previously reported as *M. tuberculosis-*dormancy biomarkers [12]. The metabolic activity marker fluorescein diacetate stained *M. ulcerans* mycobacteria either green fluorescent or not (Fig. 1, Fig. S2) [12]. However, in all the tubes incubated under ambient air, all the 6 *M. ulcerans* strains stained positively with fluorescein diacetate positive whereas and no Nile red-positive bodies were observed.

## Microcalorimetry test results

After 100 hours of heat-flow measurements by the microcalorimeter instrument, we observed a heat-flow production by dynamic *M. ulcerans* 33728, *M. ulcerans* 25894 and *M. ulcerans* 25900 strains (Fig. 2). The dynamic *M. ulcerans* 25900 strain reached a heat-flow peak of ≈2.83 µW after 36-hour incubation, while a heat-flow peak of ≈3.89 μW for *M. ulcerans* 33728 strain and ≈1.84 µW for *M. ulcerans* 25894 strain was reached after 92-hour incubation and 94-hour incubation, respectively. Dead mycobacteria used as negative controls produced no heat-flow during all the period of microcalorimetry experiment for any the three investigated strains (Fig. 2). Apart from the dormant *M. ulcerans* 33728 strain and the dormant *M. ulcerans* 25900 strain for which no heat-flow has been detected by microcalorimetry test, interestingly, one of the triplicate dormant *M. ulcerans* 25894 strain produced a heat-flow that reached the peak of 4.06 µW after ≈77 hours.

**Fig. 2.**
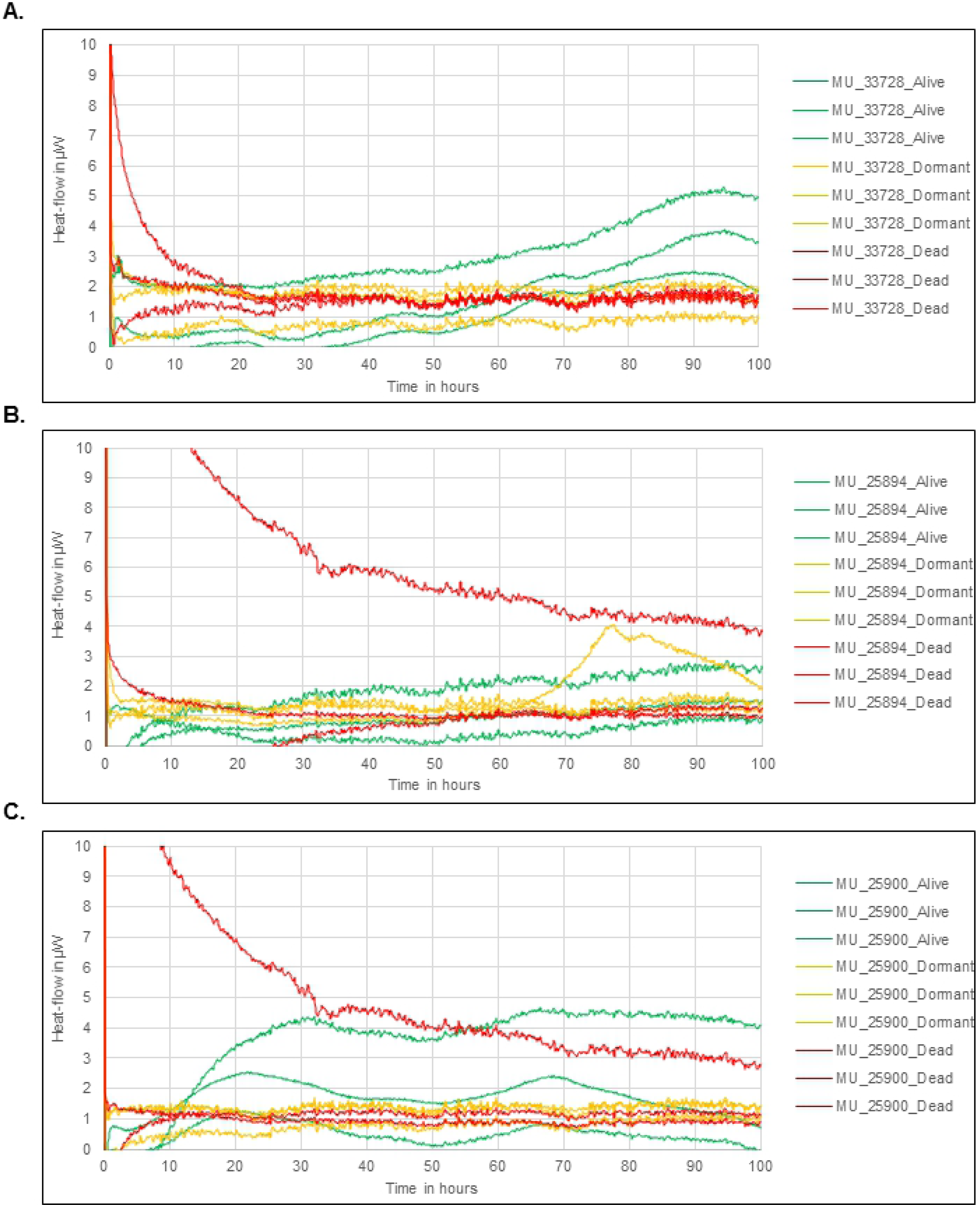
Heat-flow produced by three different *M. ulcerans* strains, according to their biological state **(A)** *M. ulcerans* 33728 strain; **(B)** *M. ulcerans* 25894 strain; **(C)** *M. ulcerans* 25900 strain. Green curves: dynamic mycobacteria; orange curves: dormant mycobacteria; red curves: dead mycobacteria.

### Subculture of dormant *M. ulcerans* strains

Two-month-old anaerobic cultures plated on Middlebrook 7H10 agar medium, grew colonies after 14-27-day incubation at 30°C under ambient air, except for *M. ulcerans* ATCC CU001 which never regrew after 6-week observation.

## DISCUSSION

Experimental data here reported indicated that *M. ulcerans* was able to sustain viability two months after progressive depletion in oxygen, with criteria for dormancy; a previously unreported property of the species *M. ulcerans*. These experimental data have been validated by the negativity of the negative controls along with congruent conclusions achieved by three experimental approaches. With respect to its phylogenetic position among mycobacteria, the fact that *M. ulcerans* exhibited dormancy was not entirely surprising: dormancy has already been reported for *Mycobacterium tuberculosis* [13], *Mycobacterium bovis* BCG [16], *Mycobacterium smegmatis* [17] and *Mycobacterium avium* subspecies *paratuberculosis* [18]. In this study, microcalorimetry was used for the first time to investigate the physiology of *M. ulcerans*. Indeed, microcalorimetry had been previously used in determining the minimum inhibitory concentration of isoniazid, ethambutol, and moxifloxacin against *M. smegmatis, M. avium*, and *M. tuberculosis* [19]; but never incorporated *M. ulcerans*, to our knowledge. Here, microcalorimetry yielded data aligned with those achieved using the DDD staining as well as culture; and indicated that metabolism of dormant *M. ulcerans* was reduced to the point that heat flow curves of dormant *M. ulcerans* were similar to those of dead *M. ulcerans*; except that one dormant *M. ulcerans* 25894 strain did produce detectable increasing heat, firmly indicated that this strain was not dead, but indeed viable and dormant.

A second, unanticipated observation was that the extent of dormancy was dependent on the *M. ulcerans* strain. In fact, one strain *M. ulcerans* CU001 died during progressive depletion in oxygen, and we have no explanation for this observation. In particular, methylene blue here used to monitor oxygen depletion in the tubes, has been used here at a concentration 100 times below the minimal inhibitory concentration we previously reported for this particular strain [20]. This observation was unexpected because strain-dependent dormancy was not previously reported for any other investigated mycobacteria.

Indeed, 38 of 48 genes reported in the dormancy operon DosR in *M. tuberculosis*, are retrievable *M. ulcerans* genomes we previously examined [21], in particular, the key regulator gene *dosR* was detected in these *M. ulcerans* genomes [21].

Whether *M. ulcerans* dormancy could be of clinical interest beyond the environmental aspect remains to be investigated as dormancy correlates with an increased resistance to otherwise effective antimycobacterial drugs, in the case of *M. tuberculosis* [16]. The DDD staining here applied to *M. ulcerans*, offers a convenient tool to further explore the clinical relevance of the experimental observations here reported.

## Supplementary figures

**Fig. S1**. Hungate tubes containing six *M. ulcerans* strains at day 0 and after 27-day incubation in progressive anaerobic atmosphere according to Wayne dormancy model. This panel features changing color of the culture medium, from blue to transparency, indicative of oxygen depletion.

**Fig. S2**. DDD staining on Wayne model of dormancy after 02-month incubation of *M. ulcerans* CU001, *M. ulcerans* ATCC 19423, *M. ulcerans* ATCC 25900, *M. ulcerans* ATCC 25894 and *M. ulcerans* ATCC 25898. Microscopic observation was performed using a 63X/1.4NA oil immersion objective of Zeiss Spinning Disk.

**Supplementary Table.**
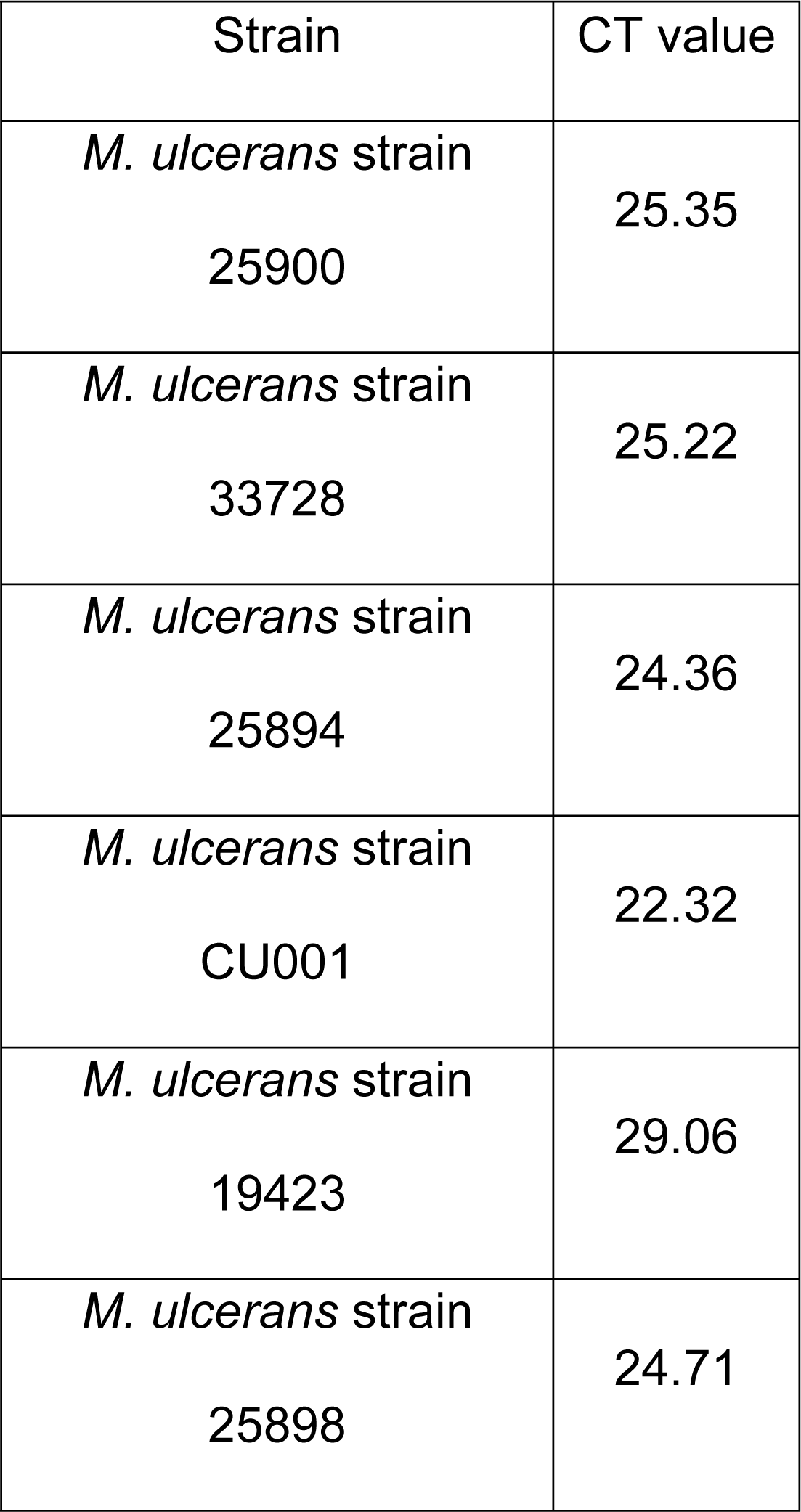
Real-time PCR cycle threshold (CT) values for *M. ulcerans* strains sub cultured after dormancy.

## REFERENCES

1. Zingue D, Bouam A, Tian RBD, Drancourt M. Buruli Ulcer, a Prototype for Ecosystem-Related Infection, Caused by *Mycobacterium ulcerans*. Clin Microbiol Rev. 2017;3: e00045–17. doi: 10.1128/CMR.00045-17.

2. George KM, Chatterjee D, Gunawardana G, Welty D, Hayman J, Lee R, et al. Mycolactone: a polyketide toxin from *Mycobacterium ulcerans* required for virulence. Science 1999;283: 854–857. doi: 10.1126/science283.5403.854

3. Mve-Obiang A, Lee RE, Portaels F, Small PL. Heterogeneity of mycolactones produced by clinical isolates of *Mycobacterium ulcerans*: implications for virulence. Infect Immun. 2003;71: 774–783. doi: 10.1128/iai.71.2.774-783.2003

4. Guenin-Macé L, Veyron-Churlet R, Thoulouze MI, Romet-Lemonne G, Hong H, Leadlay PF, et al. Mycolactone activation of Wiskott-Aldrich syndrome proteins underpins Buruli ulcer formation. J Clin Invest. 2013;123: 1501–1512. doi: 10.1172/JCI66576

5. Chany AC, Veyron-Churlet R, Tresse C, Mayau V, Casarotto V, Le Chevalier F, et al. Synthetic variants of mycolactone bind and activate Wiskott-Aldrich syndrome proteins. J Med Chem. 2014;57: 7382–7395. doi: 10.1021/jm5008819

6. Portaels F, Meyers WM, Ablordey A, Castro AG, Chemlal K, de Rijk P, et al. First cultivation and characterization of *Mycobacterium ulcerans* from the environment. PLoS Negl Trop Dis. 2008;2: e178. doi: 10.1371/journal.pntd.0000178

7. Van Zyl A, Daniel J, Wayne J, McCowan C, Malik R, Jelfs P, et al. *Mycobacterium ulcerans* infections in two horses in south-eastern Australia. Aust Vet J. 2010;88: 101–106. doi: 10.1111/j.1751-0813.2009.00544.x

8. O’Brien CR, McMillan E, Harris O, O’Brien DP, Lavender CJ, Globan M, et al. Localised *Mycobacterium ulcerans* infection in four dogs. Aust Vet J. 2011;89: 506–510. doi: 10.1111/j.1751-0813.2011.00850.x

9. Owusu E, Newman MJ, Akumwena A, Bannerman E, Pluschke G. Evaluating decontamination protocols for the isolation of *Mycobacterium ulcerans* from swabs. BMC Microbiol. 2017;1: 2. doi: 10.1186/s12866-016-0918-x

10. Käser M, Rondini S, Naegeli M, Stinear T, Portaels F, Certa U, et al. Evolution of two distinct phylogenetic lineages of the emerging human pathogen *Mycobacterium ulcerans*. BMC Evol Biol. 2007;7:177. doi: 10.1186/1471-2148-7-177

11. Hammoudi N, Saad J, Drancourt M. The diversity of mycolactone-producing mycobacteria Microb Pathog. 2020;104362. doi: 10.1016/j.micpath.2020.104362

12. Loukil A, Darriet-Giudicelli F, Eldin C, Drancourt M. Pulmonary Tuberculosis Conversion Documented by Microscopic Staining for Detection of Dynamic, Dormant, and Dead Mycobacteria (DDD Staining). J Clin Microbiol. 2018;56: e01108–18. doi: 10.1128/JCM.01108-18

13. Wayne LG, Hayes LG. An *in vitro* model for sequential study of shiftdown of *Mycobacterium tuberculosis* through two stages of nonreplicating persistence. Infect Immun. 1996;64: 2062–2069. doi: 10.1128/IAI.64.6.2062-2069.1996

14. Braissant O, Wirz D, Göpfert B, Daniels AU. Biomedical use of isothermal microcalorimeters. Sensors (Basel). 2010;10:9369–9383. doi: 10.3390/s101009369

15. Fyfe JA, Lavender CJ, Johnson PD, Globan M, Sievers A, Azuolas J, et al. Development and application of two multiplex real-time PCR assays for the detection of *Mycobacterium ulcerans* in clinical and environmental samples. Appl Environ Microbiol. 2007;73: 4733–4740. doi: 10.1128/AEM.02971-06

16. Boon C, Dick T. *Mycobacterium bovis* BCG response regulator essential for hypoxic dormancy. J Bacteriol. 2002;184: 6760–6767. doi: 10.1128/jb.184.24.6760-6767.2002

17. Mayuri BG, Das TK, Tyagi JS. Molecular analysis of the dormancy response in *Mycobacterium smegmatis*: expression analysis of genes encoding the DevR-DevS two-component system, Rv3134c and chaperone alpha-crystallin homologues. FEMS Microbiol Lett. 2002;211:231–7. doi: 10.1111/j.1574-6968.2002.tb11230.x.

18. Parrish N, Vadlamudi A, Goldberg N. Anaerobic adaptation of *Mycobacterium avium* subspecies *paratuberculosis* in vitro: similarities to *M. tuberculosis* and differential susceptibility to antibiotics. Gut Pathog. 2017;9:34. doi: 10.1186/s13099-017-0183-z

19. Howell M, Wirz D, Daniels AU, Braissant O. Application of a microcalorimetric method for determining drug susceptibility in *Mycobacterium* species. J Clin Microbiol. 2012;50: 16–20. doi: 10.1128/JCM.05556-11.

20. Tian RB, Asmar S, Napez C, Lépidi H, Drancourt M. Effectiveness of purified methylene blue in an experimental model of *Mycobacterium ulcerans* infection. Int J Antimicrob Agents. 2017;49: 290–295. doi: 10.1016/j.ijantimicag.2016.11.012

21. Chen T, He L, Deng W, Xie J. The *Mycobacterium* DosR regulon structure and diversity revealed by comparative genomic analysis. J Cell Biochem. 2013;114: 1–6. doi: 10.1002/jcb.24302

